# DCAlign v1.0: Aligning biological sequences using co-evolution models and informative priors

**DOI:** 10.1101/2022.05.18.492471

**Authors:** Anna Paola Muntoni, Andrea Pagnani

## Abstract

DCAlign is a new alignment method able to cope with the conservation and the co-evolution signals that characterize the columns of a multiple sequence alignment of homologous sequences. However, the pre-processing steps required to align a candidate sequence are computationally demanding. We show in v1.0 how to dramatically reduce the overall computing time by including an empirical prior over an informative set of variables mirroring the presence of insertions and deletions.

**Availability and implementation:** DCAlign v1.0 is implemented in Julia and it is fully available at https://github.com/infernet-h2020/DCAlign

## I. INTRODUCTION

A common task in Bioinformatics is to cast evolutionary related biological sequences into a Multiple Sequence Alignment (MSA). The objective of this task is to identify and align conserved regions of the sequences by maximizing the similarity among the columns of the MSA. State-of-the-art alignment methods, like `HMMER` [3] or `Infernal` [9] (for protein and RNA sequences respectively), use hand curated MSAs of small representative subsets of the sequences to be aligned (the *seed* alignments) to build profile Hidden Markov Models (HMM) that are used to align candidate sequences. However, homologous sequences show signals of correlated mutations (epistasys) undetected by profile models.

Conservation and co-evolution signals are at the basis of Direct Coupling Analysis (DCA)-based statistical models [2, 6]. Recently, these models have been used to perform pairwise alignments [10], remote homology search [11] and to align biological sequences to a seed model [7]. The latter method (viz. `DCAlign`) returns the ordered sub-sequence of a query unaligned sequence which maximizes an objective function related to the DCA model of the seed. Standard DCA models fail to adequately describe the statistics of insertions and gaps.

To alleviate this limitation, we introduced in the objective function gap and insertion penalties, learned from the seed alignment. While for the insertions penalty the computational complexity is negligible, inferring gap penalties is a time-consuming problem (see [7] and Supplementary Note). Here, we treat penalties in terms of empirical informative priors computed from the seed sequences. The prior parameters (both for gaps and insertions) are extracted from the seed alignment in an unsupervised manner. Finally, to further speed-up the learning of the seed-based objective function, we obtain the parameters of the DCA model using pseudo-likelihood maximization [4] instead of Boltzmann Machine Learning [5, 8]. `DCAlign`, is a computational pipeline that allow to compute the seed-model parameters in a few minutes, contrary to its original implementation which required at least a day of computation in the best scenario. The constrained optimization problem is solved approximately through a message-passing algorithm (see Supplementary Note).

## II. IMPLEMENTATION

Our algorithm estimates the optimal ordered sub-sequence compatible with a DCA model and empirical knowledge of the seed. Let ***A*** be an unaligned sequence of length *N*, and ***S*** the aligned representation of length *L*. We only consider the *L < N* case. At each *i* = 1, …, *L*, we define a Boolean variable *x*_*i*_ ∈ {0, 1} and a pointer *n*_*i*_ ∈ [0, …, *N* + 1] [7]. The variable *x*_*i*_ indicates whether the position *i* is a *gap* ‘-’ (*x*_*i*_ = 0) or a *match, i*.*e*. a symbol in ***A***. When *i* is a match, *n*_*i*_ identifies where *S*_*i*_ matches ***A***, i.e. 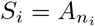; instead, for *x*_*i*_ = 0, the value of *n*_*i*_ is used for keeping track of the last *match* symbol before the *i*^th^ column. To allow this information to propagate through both gapped and matched states, we impose that *n*_*i*+1_ *> n*_*i*_ if a *match* occurs at *i* + 1, otherwise we set *n*_*i*+1_ = *n*_*i*_.

Let us define a pointer-difference variable as 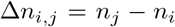 for *i* = 1, …, *L* and *j > i*. Each auxiliary variable 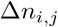 quantifies how many symbols of the unaligned sequence ***A*** are present between two *i, j* positions of the aligned counterpart ***S***. We illustrate in Fig. 1 (a) (left panel) the most relevant cases. For instance, when 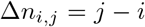, no insertion and no gap appear between *i* and *j* as the number of symbols in the aligned (*j − i*) and unaligned (*n*_*j*_ −*n*_*i*_) sequence coincide. Alternatively, 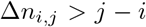 denotes the presence of insertions whereas if 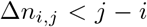 gaps are present between *i* and *j* (see Supplementary Note for an extensive discussion).

**FIG. 1.**
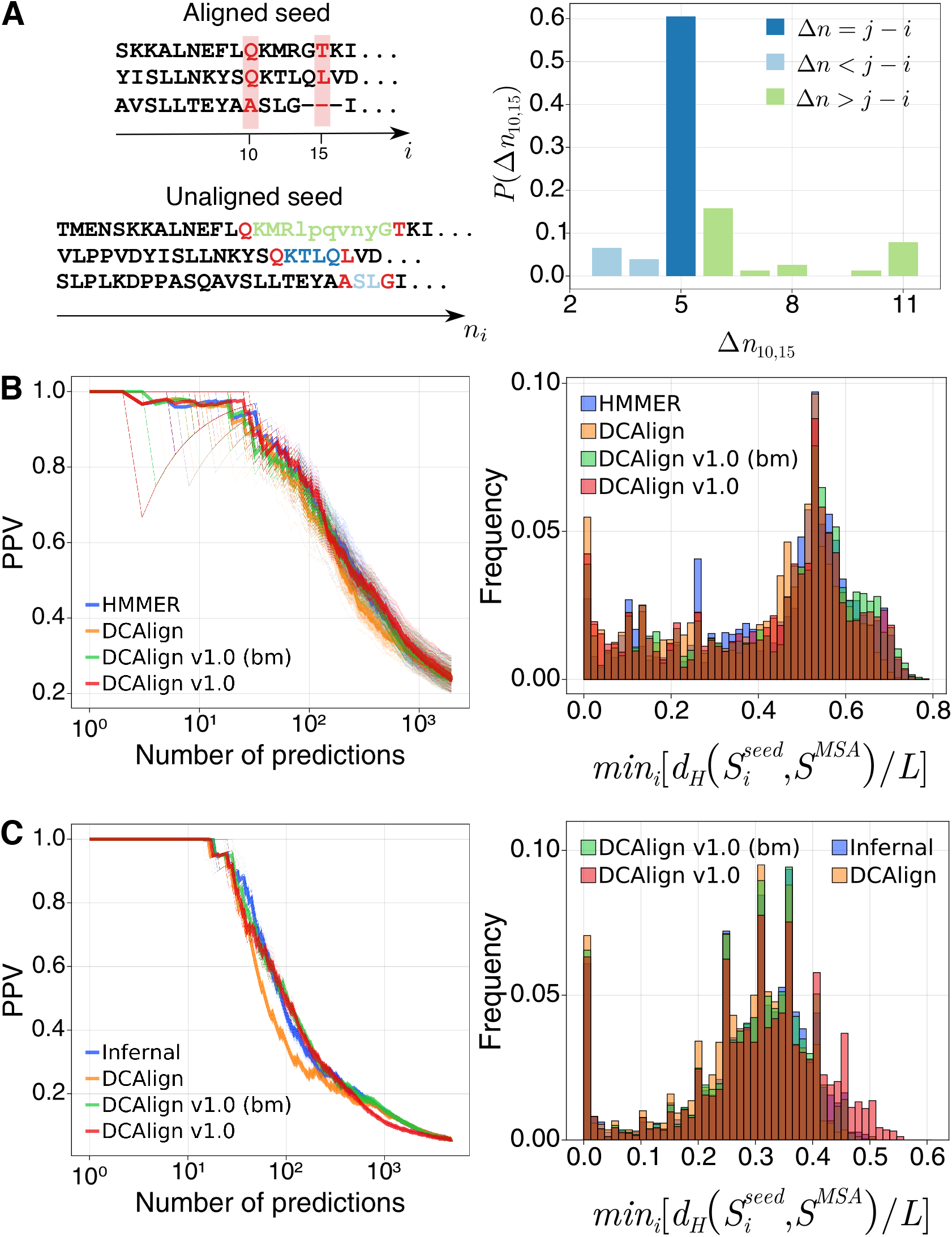
(a) Schematic representation of the pointer-differences variables for indices *i* = 10 and *j* = 15 in three seed sequences (left) and empirical probability *P* (Δ*n*_10,15_) (right) for PF00035. (b) and (c) PPV curves (the thick line shows the average behavior among different known structures) and proximity histograms for PF00035 and RF00167 respectively.

As shown in Fig. 1 (a), we can infer from the seed sequences an empirical probability 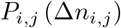 for all pairs *i, j*, by evaluating the normalized histograms of the realizations of the *n*_*j*_ − *n*_*i*_. This last step requires a few seconds of computation on an ordinary laptop.

Finally, we can express the target constrained maximization problem as

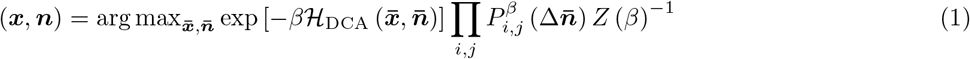

where ℋ_DCA_ is the Potts Hamiltonian, i.e. the DCA model, describing the seed, *Z* and *β* are, in the statistical mechanics jargon, the partition function and the inverse temperature parameter respectively. The maximization only runs over the feasible assignment of the variables. The informative prior will guide the optimization process towards solutions that, among those that maximize the Boltzmann distribution associated with ℋ_DCA_, reproduce the statistics of the pointer differences and, as a consequence, the (position-specific) gap and insertion statistics of the seed alignment. Unfortunately, the problem thus stated is unfeasible as the partition function cannot be efficiently computed. Similarly to the first `DCAlign` version, we resort to an approximate message-passing algorithm here coupled to an annealing scheme over *β* to get the best assignment of the variables, namely the aligned sequence (see Supplementary Note).

Once the seed model is learned, the alignment scheme alone requires the same running time needed by the first implementation of the algorithm, i.e. an average of few seconds for *N* ∼ 100.

## III. RESULTS

To understand whether the introduction of the prior affects the performances of the alignment procedure, we align the sequences of Pfam PF00035 and Rfam RF00167 using (i) a state-of-the-art method HMMER or Infernal for PF00035 or RF00167 respectively (ii) `DCAlign`, the implementation presented in [7], (iii) `DCAlign v1.0` (bm) which differs from `DCAlign` only for the presence (absence) of the informative prior (extra penalties) (iv) `DCAlign v1.0` which differs from `DCAlign v1.0 (bm)` for the ℋ_DCA_ learned by pseudo-likelihood maximization instead of Boltzmann machine learning. The seed of the two families contains a few sequences (76 and 133 for PF00035 and RF00167 respectively) therefore in `DCAlign` the gap penalties have required a long learning step of several days. We quantitatively compare the performances of all the methods by evaluating a DCA model associated with each method-based alignment and by a proximity comparison between aligned and seed sequences. Figs. 1 (b) and (c) show the performances of all the methods for PF00035 and RF00167 respectively in terms of the positive predictive value (PPV) curve associated with the prediction of contacts performed by the DCA models for several known structures of the Protein Data Bank [1] (left panels) and a histogram of the Hamming distances of the closest seed sequences to the aligned ones (right panels) (see Supplementary Note for more details). Both the PPV curves and the proximity histograms quite overlap for PF00035 and RF00167 suggesting that `DCAlign v1.0` reaches comparable performances to the original implementation and state-of-the-art methods.

## IV. CONCLUSION

`DCAlign v1.0` is a new implementation of the DCA-based alignment technique, `DCAlign`, which conversely to the first implementation, allows for a fast parametrization of the seed alignment. The new modeling significantly drops the pre-processing times and guarantees a qualitatively equivalent alignment of a set of target sequences.

## ACKNOWLEDGEMENTS

APM and AP acknowledge financial support from Marie Sklodowska-Curie, grant agreement no. 734439(INFERNET). We also warmly thank Indaco Biazzo, Alfredo Braunstein, Louise Budzynski and Luca Dall’Asta for interesting discussions.

## DATA AVAILABILITY

`DCAlign v1.0` is available at https://github.com/infernet-h2020/DCAlign

## Notes

### Competing Interest Statement

The authors have declared no competing interest.

