## Supplemental Material for "DCAlign v1.0: Aligning biological sequences using co-evolution models and informative priors"

### Supplementary data for
### DCAlign v1.0: Aligning biological sequences using co-evolution models and informative priors

Anna Paola Muntoni[1] and Andrea Pagnani[1, 2, 3]
[1]*Italian Institute for Genomic Medicine, IRCCS Candiolo, SP-142, I-10060 Candiolo (TO), Italy*
[2]*Politecnico di Torino, Corso Duca degli Abruzzi, 24, I-10129, Torino, Italy*
[3]*INFN, Sezione di Torino, Torino, Italy*

#### CONTENTS

I. Modeling alignments 1
II. Standard learning of the gap penalties 2
III. Empirical prior over the pointer-difference variables 2
IV. Approximate message-passing equations 3
V. Annealing scheme 4
VI. Computation of the Positive Predicted Value curves 5
VII. Computation of the proximity histograms 5
VIII. Learning the co-evolution model using pseudo-likelihood maximization 6
References 6

#### I. MODELING ALIGNMENTS

In the following we briefly summarize how to model aligned sequences according to `DCAlign`. Given the notation introduced in the main text, to obtain an ordered sequence suffices to impose *local* constraints on the variables $(\boldsymbol{x}, \boldsymbol{n})$ through a compatibility function $\chi_{i-1,i}\left(x_{i-1}, n_{i-1}, x_i, n_i\right)$ equals to one only for feasible assignments, namely

$$\begin{aligned}
\chi_{i-1,i}\left(0, n_{i-1}, 0, n_i\right) &= \mathbb{I}\left[n_i = n_{i-1}\right] \\
\chi_{i-1,i}\left(1, n_{i-1}, 0, n_i\right) &= \mathbb{I}\left[n_i = n_{i-1} \vee n_i = N+1\right] \\
\chi_{i-1,i}\left(0, n_{i-1}, 1, n_i\right) &= \mathbb{I}\left[0 \le n_{i-1} < n_i < N+1\right] \\
\chi_{i-1,i}\left(1, n_{i-1}, 1, n_i\right) &= \mathbb{I}\left[0 < n_{i-1} < n_i < N+1\right]
\end{aligned} \tag{1}$$

where $\mathbb{I}\left[\mathcal{E}\right]$ is the indicator function of event $\mathcal{E}$. The original problem can be cast into a constrained optimization problem over the Boolean and pointer variables as

$$(\boldsymbol{x}, \boldsymbol{n}) = \arg\max_{(\bar{\boldsymbol{x}}, \bar{\boldsymbol{n}})} Z\left(\beta\right)^{-1} e^{-\beta \tilde{\mathcal{H}}(\bar{\boldsymbol{x}}, \bar{\boldsymbol{n}})} \prod_{i,i+1} \chi_{i,i+1}\left(\bar{x}_i, \bar{n}_i, \bar{x}_{i+1}, \bar{n}_{i+1}\right) \tag{2}$$

where $Z$ is the partition function, i.e. the normalization of the joint distribution of the variables

$$Z\left(\beta\right) = \sum_{\substack{\boldsymbol{x}, \boldsymbol{n}: \\ \prod_{i,i+1} \chi_{i,i+1} = 1}} e^{-\beta \tilde{\mathcal{H}}(\boldsymbol{x}, \boldsymbol{n})} \tag{3}$$

and $\tilde{\mathcal{H}}$ is (minus) the objective function, namely the Hamiltonian characterizing the seed

$$\tilde{\mathcal{H}}\left(\boldsymbol{x}, \boldsymbol{n}\right) = \mathcal{H}_{\text{DCA}}\left(\boldsymbol{x}, \boldsymbol{n}\right) - \mathcal{H}_{\text{ins}}\left(\boldsymbol{x}, \boldsymbol{n}\right) - \mathcal{H}_{\text{gap}}\left(\boldsymbol{x}, \boldsymbol{n}\right) \tag{4}$$

The term $\mathcal{H}_{\text{DCA}}(\boldsymbol{x}, \boldsymbol{n})$ is the Potts Hamiltonian learned from the seed sequences using standard inverse modeling procedures

$$\mathcal{H}_{\text{DCA}}(\boldsymbol{x}, \boldsymbol{n}) = -\sum_{i,j} J_{i,j}\left(A_{x_i n_i}, A_{x_j n_j}\right) - \sum_i h_i\left(A_{x_i n_i}\right) \tag{5}$$

where $A_0 = `-`$. More precisely, to infer the coupling matrices and the fields, we employ Boltzmann machine learning [4] for the algorithms denoted as `DCAlign`, `DCAlign v1.0 (bm)` while for the new implementation encoded into `DCAlign v1.0` we use pseudo-likelihood maximization [2].

The other two energy terms $\mathcal{H}_{\text{ins}}$ and $\mathcal{H}_{\text{gap}}$ penalize the presence of insertions or gaps within the aligned sequences. They take the form of

$$\mathcal{H}_{\text{ins}}(\boldsymbol{x}, \boldsymbol{n}) = \sum_i \left(1 - \delta_{\Delta n_i, 0}\right)\left[\lambda_o^i + \lambda_e^i\left(\Delta n_i - 1\right)\right] \tag{6}$$

$$\mathcal{H}_{\text{gap}}(\boldsymbol{x}, \boldsymbol{n}) = \sum_i \delta_{x_i,0}\{\mu^{\text{int}}\mathbb{I}\left[0 < n_i < N+1\right] + \mu^{\text{ext}}\mathbb{I}\left[n_i = 0 \vee n_i = N+1\right]\} \tag{7}$$

where $\Delta n_i$ is the number of insertions between positions $i$ and $i-1$, $\lambda_o^i$ ($\lambda_e^i$) is the penalty for creating (adding) an insertion at position $i$ and $\mu^{\text{ext}}$ ($\mu^{\text{int}}$) is cost of using a gap symbol at the beginning or the end of the align sequence (within two symbols). While the parameters $\{\boldsymbol{\lambda}_o, \boldsymbol{\lambda}_e\}$ can be learned by means of a (fast) likelihood maximization, the learning of the gap penalties $\mu^{\text{ext}}$ and $\mu^{\text{int}}$ are determined in a supervised and slow manner (see Sec. II). To significantly decrease the computation time necessary to determine $\mathcal{H}_{\text{ins}}$ and $\mathcal{H}_{\text{gap}}$, we introduce in Sec. III an empirical prior which accommodates both the presence of insertions and gaps.

#### II. STANDARD LEARNING OF THE GAP PENALTIES

As discussed in [3], if the seed alignment contains a large number of sequences ($\geq 10^3$), it suffices to re-align all of them for each set of candidate gap penalties $\left(\mu^{\text{int}}, \mu^{\text{ext}}\right)$ and to choose those that minimize the average Hamming distance between the original seed sequences and the re-aligned ones. The trial number of gap penalties is often about 80, for $\mu^{\text{int, ext}} \in [0.0, 4.0]$ which coincides with the number of times the seed needs to be re-aligned. Considering that the computation time is on average of few seconds for a sequence of length $N$ and the number of sequences is about one hundred, the overall time is of one day.

Instead, if the seed is composed of a few sequences (i.e. less than $10^3$), the average Hamming distance is always zero for all candidate gap penalties, i.e. the re-aligned sequences always coincide with the true sequences, carrying no information about the best set of parameters. To resort to this issue, it is reasonable to introduce a validation set of sequences to be aligned using all candidate gap penalties (usually of the order of $10^3$). Alternatively to the Hamming distance, one may consider the symmetric Kullback-Leibler divergence sDKL between a seed model $\mathcal{H}_{\text{DCA}}$ and a model learned from each of the candidate aligned test set obtained for a couple $\left(\mu^{\text{ext}}, \mu^{\text{int}}\right)$, that is $\mathcal{H}_{\text{DCA}}^{\left(\mu^{\text{ext}}, \mu^{\text{int}}\right)}$, defined as

$$\text{sDKL} = \left\langle \mathcal{H}_{\text{DCA}} - \mathcal{H}_{\text{DCA}}^{\left(\mu^{\text{ext}}, \mu^{\text{int}}\right)} \right\rangle_{\mathcal{H}_{\text{DCA}}^{\left(\mu^{\text{ext}}, \mu^{\text{int}}\right)}} + \left\langle \mathcal{H}_{\text{DCA}}^{\left(\mu^{\text{ext}}, \mu^{\text{int}}\right)} - \mathcal{H}_{\text{DCA}} \right\rangle_{\mathcal{H}_{\text{DCA}}} \tag{8}$$

Here the symbol $\langle a \rangle_b$ denotes the average value of the quantity $a$ according to the Boltzmann distribution having Hamiltonian $b$. This last step not only requires to align a huge number of sequences (as for the copious seed case), but the computation of the Kullback-Leibler distance requires learning as many models as the number of candidate gap penalties, and sampling from them (to compute the expectation values) using Monte Carlo Markov Chain.

Considering that, using Boltzmann machine learning, the learning of a DCA model costs several hours of computation, the overall process necessitates of a computational time that vary from one day to several days.

#### III. EMPIRICAL PRIOR OVER THE POINTER-DIFFERENCE VARIABLES

A much faster strategy aimed at modeling the presence of gaps and insertions exploits the information over the realization of the pointer variables $\boldsymbol{n}$ in the seed sequences.

As described in the main text, for each couple of indices of the columns of the MSA, let us denote them as $i$ and $j > i$, we compute the empirical frequencies (over the seed sequences) of the corresponding $\Delta n_{i,j} = n_j - n_i$. This

variable quantifies the number of symbols that are present, within the unaligned seed sequence, between the positions $i$ and $j$ of the aligned counterpart. For sake of clarity, let us consider the $j = i+1$ case; here $j - i = 1$ and depending on the value $\Delta n_{i,i+1}$ we can identify three scenarios. $\Delta n_{i,i+1} = 1 = j - i$ reveals that no insertion and no gap are present between $i$ and $i+1$ because the symbols appearing in the aligned sequence are consecutive in the unaligned one. Instead, if $\Delta n_{i,i+1} > 1$ more than one symbol lies between $A_{n_i}$ and $A_{n_{i+1}}$ and, in this case, $\Delta n_{i,i+1} - 1$ quantifies the number of insertions between $i$ and $i+1$. Conversely, if $\Delta n_{i,i+1} = 0 < j - i$ the two pointers $n_i$ and $n_{i+1}$ coincide and this is only possible when a gap appears in $A_{n_{i+1}}$. For a generic case $j > i+1$ the two situations may overlap meaning that both insertions and gaps can be present. However, a complete scan over all possible couples of indices allows one to discriminate between all possible cases. Therefore, studying the statistics of these pointer-difference variables reveals how likely is, on average, to find a certain number of insertions and gaps for a given couple of MSA positions. Independently of the nature of the possible discrepancy between $\Delta n_{i,j}$ and $i - j$, adding an informative empirical prior to the objective function informs the pointer variables to be inferred about their statistical behavior within the seed sequences.
Let us define $\Lambda_{i,j}(\Delta n)$ the normalized histogram of the pointer-difference variables associated with columns $i$ and $j$. Note that the number of possible argument $\Delta n$ of the empirical probability $\Lambda_{i,j}(\Delta n)$ depends of the length of the longest sequence of the seed. Before adding the empirical probability to the objective function in Eq. 2, we need to ensure that, once we align a sequence of length $N$, the maximum possible value $\Delta n = N + 1 - 0$ is covered by the empirical prior. Similarly, being $\Lambda(\Delta n)$ obtained from (usually) a few sequences, some argument may have exactly zero probability. This would prevent several assignments that may be, although improbable, the best choice for the target pointer variables. For this reason, we re-weight the empirical statistics adding a small pseudo-count equals to $1/M_{seed}$ for the unobserved states (where $M_{seed}$ is the number of seed sequences). The probability thus obtained is

$$P_{i,j}(\Delta n) \propto \left(1 - \frac{1}{M_{seed}}\right) \Lambda_{i,j}(\Delta n) + \frac{1}{M_{seed}}, \qquad \Delta n \in [0, \ldots, N+1] \tag{9}$$

where the proportionality symbol accounts for the missing normalization ensuring that $\sum_{\Delta n} P_{i,j}(\Delta n) = 1$. The alignment problem is therefore re-phrased in `DCAlign v1.0` as

$$(\boldsymbol{x}, \boldsymbol{n}) = \arg\max_{(\bar{\boldsymbol{x}}, \bar{\boldsymbol{n}})} Z(\beta)^{-1} e^{-\beta \mathcal{H}_{\text{DCA}}(\bar{\boldsymbol{x}}, \bar{\boldsymbol{n}})} \prod_{\substack{i,j: \\ j>i}} P_{i,j}^{\beta}(\Delta \bar{\boldsymbol{n}}) \prod_{i,i+1} \chi(\bar{x}_i, \bar{n}_i, \bar{x}_{i+1}, \bar{n}_{i+1}) \tag{10}$$

#### IV. APPROXIMATE MESSAGE-PASSING EQUATIONS

A brute-force solution to the optimization problem in Eq. 10 would require to compute the partition function

$$Z(\beta) = \sum_{\substack{\boldsymbol{x}, \boldsymbol{n}: \\ \prod_{i,i+1} \chi_{i,i+1} = 1}} e^{-\beta \mathcal{H}_{\text{DCA}}(\boldsymbol{x}, \boldsymbol{n})} \prod_{\substack{i,j: \\ j>i}} P_{i,j}^{\beta}(\Delta \boldsymbol{n}) \tag{11}$$

which is a hard and unfeasible computation as the number of terms involved in the sum scales as $\mathcal{O}\left((N+2)^L\right)$. In `DCAlign`, an approximation of the marginal probability densities $m_i(x_i, n_i)$ of the joint distribution of the variables (whose normalization appears in Eq. 3) are retrieved by iterating a set of simplified message-passing equations. From them, an assignment of the variables, and therefore the aligned sequence, is determined by looking at the arguments that maximize the marginal probabilities

$$(x_i, n_i) = \arg\max_{x,n} m_i(x, n) \qquad i = 1, \cdot, L. \tag{12}$$

In cases where the assignment does not satisfy the hard constraints encoded in the $\chi$-functions, it is possible to resort to a nucleation algorithm (see [3]).

Similarly to the first implementation, in `v1.0`, the general update scheme can be expressed using a transfer-matrix formulation (see Table I). The introduction of the empirical prior slightly modifies the *forward*, *backward* and *central*

| $i=1$ | $i=2,\cdots,L-1$ | $i=L$ |
|---|---|---|
| $m_1=\frac{1}{z_1}\mathcal{C}_1\mathcal{B}_1$ | $m_i=\frac{1}{z_i}\mathcal{C}_i\mathcal{F}_i\mathcal{B}_i$ | $m_L=\frac{1}{z_L}\mathcal{C}_L\mathcal{F}_L$ |
| $F_1=\frac{1}{f_1}\mathcal{C}_1$ | $F_i=\frac{1}{f_i}\mathcal{C}_i\mathcal{F}_i$ | — |
| — | $B_i=\frac{1}{b_i}\mathcal{C}_i\mathcal{B}_i$ | $B_L=\frac{1}{b_L}\mathcal{C}_L$ |

TABLE I. Sketch of the message-passing update equations. The terms $\mathcal{F}_i$, $\mathcal{B}_i$, $\mathcal{C}_i$ are given in Eqs. 13, 14 and 15.

terms as

$$\begin{aligned}\mathcal{F}_i\left(x_i,n_i\right) &= \sum_{x_{i-1},n_{i-1}} F_{i-1}\left(x_{i-1},n_{i-1}\right)e^{\beta J_{i-1,i}\left(A_{x_{i-1}n_{i-1}},A_{x_in_i}\right)}\times \\ &\quad \times\chi_{i-1,i}\left(x_{i-1},n_{i-1},x_i,n_i\right)P^{\beta}_{i-1,i}\left(n_i-n_{i-1}\right) \end{aligned} \tag{13}$$

$$\begin{aligned}\mathcal{B}_i\left(x_i,n_i\right) &= \sum_{x_{i+1},n_{i+1}} B_{i+1}\left(x_{i+1},n_{i+1}\right)e^{\beta J_{i+1,i}\left(A_{x_in_i},A_{x_{i+1}n_{i+1}}\right)}\times \\ &\quad \times\chi_{i,i+1}\left(x_i,n_i,x_{i+1},n_{i+1}\right)P^{\beta}_{i,i+1}\left(n_{i+1}-n_i\right) \end{aligned} \tag{14}$$

$$\begin{aligned}\mathcal{C}_i\left(x_i,n_i\right) &= \exp\left[\beta h_i\left(A_{x_in_i}\right)+\right. \\ &\quad +\sum_{j<i-1}\sum_{x_j,n_j}\chi_{lr}\left(x_i,n_i,x_j,n_j\right)\beta J_{i,j}\left(A_{x_in_i},A_{x_jn_j}\right)m_j\left(x_j,n_j\right)P^{\beta}_{i,j}\left(\Delta n_{i,j}\right)+ \\ &\quad \left.+\sum_{j>i+1}\sum_{x_j,n_j}\chi_{lr}\left(x_i,n_i,x_j,n_j\right)\beta J_{i,j}\left(A_{x_in_i},A_{x_jn_j}\right)m_j\left(x_j,n_j\right)P^{\beta}_{j,i}\left(\Delta n_{j,i}\right)\right] \end{aligned} \tag{15}$$

The $\chi_{lr}$ are redundant 'long-range' compatibility functions involving non-neighbor sites

$$\begin{aligned}\chi_{lr}\left(x_i,n_i,x_j,n_j\right) &= \mathbb{I}\left[i>j+1\right]\left\{\delta_{x_i,0}\mathbb{I}\left[n_i\geq n_j\right]+\delta_{x_i,1}\mathbb{I}\left[n_i>n_j\right]\right\}+ \\ &\quad \mathbb{I}\left[i<j-1\right]\left\{\delta_{x_j,0}\mathbb{I}\left[n_i\leq n_j\right]+\delta_{x_j,1}\mathbb{I}\left[n_i<n_j\right]\right\} \end{aligned} \tag{16}$$

derived from the short-range ones. Eqs. 13, 14 and 15 may be iteratively updated, together with those in Table I, by setting the inverse temperature $\beta=1$, up to a numerical convergence. This temperature could be understood as the 'natural' temperature of the system fully determined by the inverse methods used to parametrize the seed sequences. In `DCAlign` this scheme was sufficient for obtaining concentrated marginals and therefore a straightforward decoding to get the aligned sequence. However, in `v1.0`, at numerical convergence, very often the marginal probabilities $m_i\left(x_i,n_i\right)$ are dispersed over several states signaling a coexistence of energetically equivalent alignments, at $\beta=1$. This renders the final assignment of the variables hard, as the naif maximization of the marginal probabilities may lead to an unfeasible assignment (see Fig. 1 panel A). To solve this issue, and to actually implement the maximization step in Eq. 10, we resort to an annealing scheme over $\beta$.

#### V. ANNEALING SCHEME

To allow the message-passing dynamics to polarize on (one of) the best possible assignment of the variables, we increase the inverse temperature $\beta$ within the iterations of the equations, i.e. each $\Delta t=10$ iterations $\beta$ is increased of $\Delta\beta=0.05$. We check the polarization of the marginals by looking at the minimum value, among the sites, of the maximum value, over the states, of the $m_i\left(x_i,n_i\right)$. When this number is larger than a threshold and the assignment retrieved from the arguments that maximize the marginal probabilities is feasible, we stop the algorithm. Empirically we observe that a threshold equals to 0.30 suffices to guarantee polarized and feasible assignment. An example is reported in Fig. 1 for a sequence belonging to Pfam PF00035. We show in panels A the results for $\beta=1$ while in panel B we display the marginals obtained through the annealing scheme. The left (right) plot shows the conditional probability for each site $i$ (in the rows of the matrix) to be aligned to position $n_i$ (in the columns) for $x_i=1$ (for $x_i=0$). In Fig. 1 panel A the marginal probability is dispersed over several values of $n_i$ for $i<16$. In particular, the assignment provided by the maximization of the marginals is unfeasible because the sites $i=14,15$ both point to $n=33$. The issue is solved using the annealing over $\beta$ as displayed in panel B: site $i=14$ is now pointing to $n_{14}=33$ for $x_{14}=1$ while in $i=15$ a gap appears.

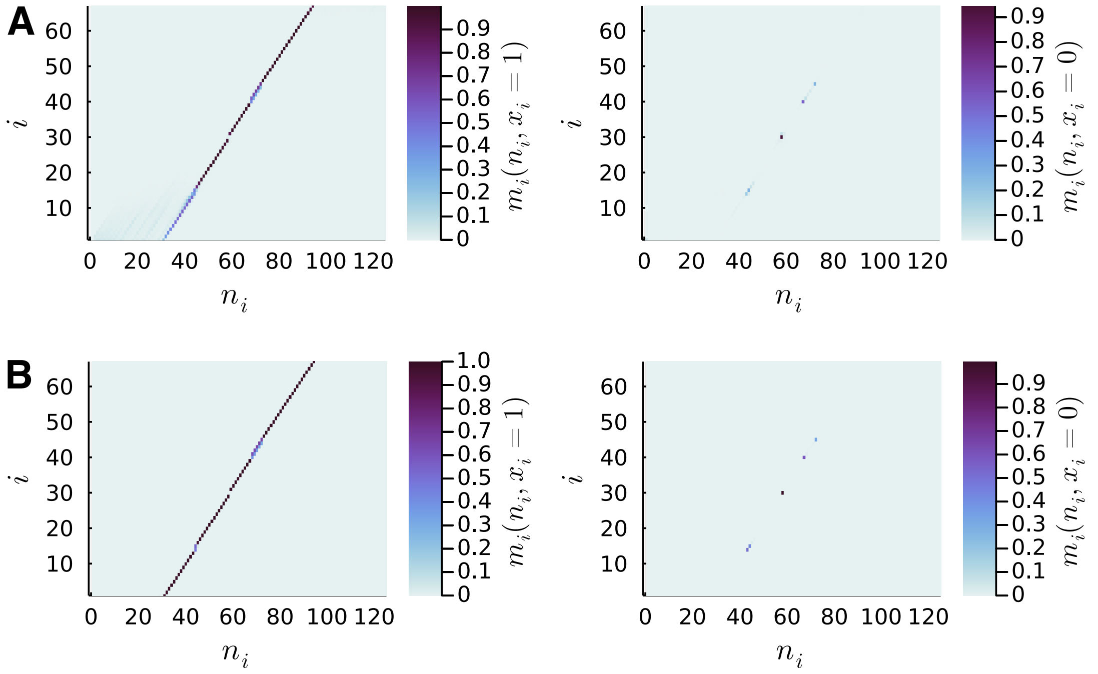


FIG. 1. Panel A shows the marginal probabilities $m_i\left(x_i = 1, n_i\right)$ (right panel) and $m_i\left(x_i = 0, n_i\right)$ (left panel) for all the sites $i = 1, \cdot, L$ when the message-passing equations are iterated at $\beta = 1$. Panel B shows instead how they slightly change when we perform the annealing over $\beta$. Probabilities for $i < 16$ appear concentrated in panel B rendering the assignment step straightforward. The target protein domain belongs to PF00035 and appears within the sequence `A0A2U9BW26_SCOMX`

#### VI. COMPUTATION OF THE POSITIVE PREDICTED VALUE CURVES

To verify whether the introduction of the empirical prior affects the quality of the alignment, we compare the outcomes provided by `v1.0` to those obtained by `DCAlign` and state-of-the-art techniques. The idea is to infer a DCA model (learned using pseudo-likelihood maximization) for each candidate multiple sequence alignment whose parameters are used to perform a contact prediction of the associated domain. In particular, for each DCA model, we compute the set of average product corrected Frobenius norms of the coupling matrices [2] which quantify how likely the pairs of sites are in contact according to the model.

To evaluate the quality of the prediction, we use as ground-truths several experimentally obtained structures downloaded from the Protein Data Bank [1] (90 for PF00035 and 46 for RF00167). Note that to allow for any comparison, we first need to map the residue sites of the crystallized sequence to those of the multiple sequence alignment. In principle, this mapping may change depending on the used alignment tool. To unify the mapping we align the sequence of the crystallized domain using the Hidden Markov Model associated with the seed sequences. Then, using the sorted Frobenius norms as scores, we compute the Positive Predictive Value (PPV) curve for each method and for each PDB entry. Thin lines in Fig. 1 of the main text are associated with each available structure, while the thick line is an average PPV curve evaluated over the different PDB structures. The plots suggest, by considering both single-entry and average comparisons, that all methods perform comparable contact structure predictions.

#### VII. COMPUTATION OF THE PROXIMITY HISTOGRAMS

Together with the contact prediction we use a sequence-based metric to evaluate the quality of the new alignment scheme. The measure we consider consists of the distribution of the proximity distances between the seed and the multiple sequence alignments obtained for the considered methods. Let us define as $\mathbf{S}^{seed}$ the set of the seed sequences and $\mathbf{S}^{MSA}$ the query sequences aligned by of the considered methods. For each sequence $S_j^{MSA}$ we compute the Hamming distance $d_H$ between $S_j^{MSA}$ and all the seed sequences and we keep the minimum distance

$$d_j = min_i d_H\left(S_j^{MSA}, S_i^{seed}\right) \tag{17}$$

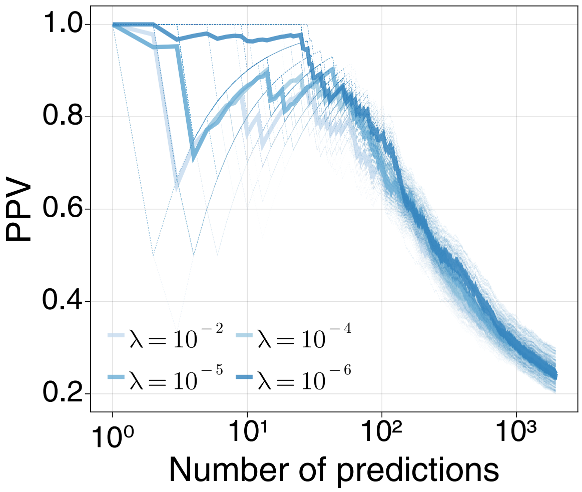


FIG. 2. PPV curves associated with four multiple sequence alignments obtained by `v1.0` for 90 known structures of PF00035 (thin dashed line are associated with the 90 considered PDB entries while thick line denotes the average behavior). The MSAs differ on the regularization strength $\lambda$ used within the pseudo-likelihood maximization scheme necessary to learn a seed model.

Then, we plot in Fig. 1 the frequencies of the minimal distances $\mathbf{d}$ achieved by each method. It is worth noting that the histograms quite overlap suggesting that the introduction of the empirical prior does not significantly affect the proximity distance between the seed and the multiple sequence alignments obtained by `DCAlign` and `v1.0` which are concurrently similar to those obtained by `HMMER` and `Infernal` (see panel B, right plot and panel C, right plot).

#### VIII. LEARNING THE CO-EVOLUTION MODEL USING PSEUDO-LIKELIHOOD MAXIMIZATION

Conversely to the first implementation of `DCAlign`, we propose in `v1.0` to learn the co-evolution model using `PlmDCA` [2]. For copious seed, the usual learning (i.e. when using default parameters) suffices to obtain a model which well align a test sequence, whereas for seed containing few sequences, we obtain better results (comparable to those obtained using a Boltzmann machine) when decreasing the regularization strength, that is when using $\lambda_J = \lambda_h = \lambda = 10^{-6}$. In Fig. 2 we plot the PPV curves associated with the DCA models learned from several multiple sequence alignments. Each MSA was obtained using a DCA model for the seed learned by setting $\lambda = \{10^{-2}, 10^{-4}, 10^{-5}, 10^{-6}\}$ (the intensity of the color of the curves mirrors the value of the regularization, the darker the color the smaller the $\lambda$ used). Thin dashed lines represent the prediction of one of the considered 90 structures while thick lines display the average behavior among the different structures. The plot suggests that decreasing the regularization strength allows us to obtain a seed model that, if used within the alignment process, guarantees an alignment of the query sequences that better mirrors the structure of the associated protein domain.

[1] Berman, H. M., Westbrook, J., Feng, Z., Gilliland, G., Bhat, T. N., Weissig, H., Shindyalov, I. N., and Bourne, P. E. (2000). The Protein Data Bank. *Nucleic Acids Research*, **28**(1), 235–242.

[2] Ekeberg, M., Lövkvist, C., Lan, Y., Weigt, M., and Aurell, E. (2013). Improved contact prediction in proteins: Using pseudolikelihoods to infer Potts models. *Physical Review E*, **87**(1), 012707. Publisher: American Physical Society.

[3] Muntoni, A. P., Pagnani, A., Weigt, M., and Zamponi, F. (2020). Aligning biological sequences by exploiting residue conservation and coevolution. *Physical Review E*, **102**(6), 062409. Publisher: American Physical Society.

[4] Muntoni, A. P., Pagnani, A., Weigt, M., and Zamponi, F. (2021). adabmDCA: adaptive Boltzmann machine learning for biological sequences. *BMC Bioinformatics*, **22**(1), 528.
